# Anatomical 3D Reconstruction of Murine Lymph Nodes for Visualization, Quantitation, and Numerical Simulation

**DOI:** 10.1101/2025.11.25.690560

**Authors:** Michael J. Donzanti, Yasaman Moghadamnia, Ashlyn T. Kapinski, Ryan Zurakowski, Jason P. Gleghorn

## Abstract

Lymph nodes (LNs) function as pharmacological sanctuary sites in HIV and metastatic cancer due to anatomical barriers that limit drug penetration. Accurate 3D reconstructions of lymph node architecture are essential for computational modeling of drug transport, yet existing methods lack compartment-specific resolution, accessibility, or throughput. Here, we present a scalable, high-fidelity pipeline for the 3D anatomical reconstruction of murine LNs, integrating optimized vibratome sectioning, multiplexed immunofluorescence staining, confocal microscopy, and custom automated segmentation algorithms. Our method precisely reconstructs key LN compartments, including lobules and high endothelial venules, with high spatial accuracy, achieving Sørensen-Dice indices >0.93 for lobules and contour-matching scores up to 77% for vasculature, as validated by quantitative comparison to manual segmentation. Compared to existing methodologies, this pipeline markedly reduces reagent usage (∼ 88%), labor time (∼ 97%), and technical complexity, offering a broadly accessible and efficient approach to high-fidelity 3D anatomical reconstruction of LN architecture. These digital twins can support computational simulations of drug distribution, immune cell trafficking, and spatial pharmacokinetics, providing critical insights into LN-resident disease mechanisms and informing therapeutic design.

## Introduction

Conventional two-dimensional (2D) imaging techniques, while accessible and cost-effective, fail to capture the inherent spatial complexity required for understanding structure-function relationships, particularly in heterogeneous and dynamic systems (1–6). Understanding the three-dimensional (3D) architecture of biological tissues is critical for accurately understanding complex physiological phenomena such as organ morphogenesis, immune cell dynamics, and therapeutic drug transport (7–10). Additionally, it is well appreciated that tissue geometry is dynamic and can remodel in response to acute and chronic stimuli, which can significantly alter tissue function and potential treatments (11, 12). Computational models, including digital twins, increasingly complement experimental approaches by simulating transport phenomena and cellular dynamics (13–15), but require anatomically accurate 3D geometries as input domains. By developing faster methods for 3D anatomical reconstruction, we can bridge the gap between traditional experimentation and computational methods, thereby enhancing our understanding of complex physiology. This, in turn, will lead to therapeutic design principles and dosing strategies that are applicable across populations, paving the way for precision medicine that optimizes drug efficacy and delivery.

The lymph node (LN), a highly compartmentalized (**Fig. 1A**) and immunologically important organ, exemplifies these challenges due to its small size and structural complexity, heterogeneity, and dynamic remodeling capabilities (16, 17). Specialized endothelial barriers, notably high endothelial venules (HEVs) and sinus-parenchyma interfaces, compartmentalize the LN into distinct functional regions (**Fig. 1B,C**), critically influencing immune responses and drug transport (18–25). The parenchyma of the lymph node, called the lobule, contains the cortex, paracortex, and follicles, with the LN architecture having extensive heterogeneity, exhibiting multiscale variability within a single lymph node, within a single individual, and between individuals (26–32). Diseases such as HIV and metastatic cancer exploit these anatomical complexities, as limited drug penetration into LN lobules enables disease persistence, making LNs pharmacological sanctuary sites. Despite the clinical significance, detailed quantitative understanding of how compartment-specific geometries impact molecular and cellular transport remains limited, particularly in the context of dynamic plasticity of the LN architecture in inflammation and disease (33, 34). The ability to accurately produce 3D anatomical reconstructions of the lymph node can provide information to quantify morphological characteristics and computationally map spatial drug distribution, resulting in a better understanding of how lymph node anatomy affects drug transport and, therefore, treatment efficacy for diseases such as HIV or metastatic cancer.

**Fig. 1.**
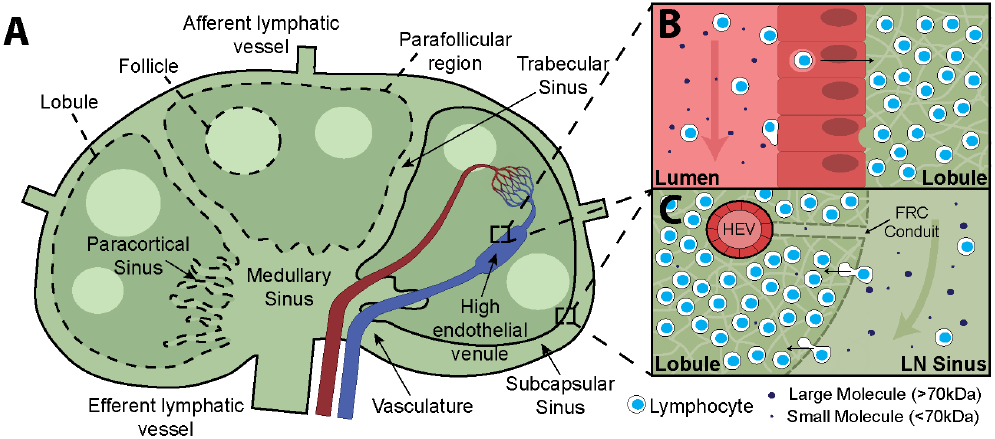
Schematic of lymph node compartmentalization. **A)** Lymph node anatomy, showing boundaries between the lobule and the **B)** blood and the **C)** sinuses. While the maintenance of these specialized endothelial barriers results in isolation for proper immune function, it also results in poor drug penetration. Most molecules of all sizes are restricted from passive diffusion across the blood-lobule barrier and most large molecules are restricted from passive diffusion across the sinus-lobule barrier.

Current imaging methods for 3D anatomical reconstruction, including micro-computed tomography (µCT), magnetic resonance imaging (MRI), serial histological sectioning, and whole-mount imaging, each present significant limitations. Volumetric imaging modalities (35–39) offer *in situ* imaging with faster image acquisition times, particularly for large whole organ scanning, but are limited in multiplexed compartment labeling and tissue scale resolution. Whereas more common histological and immunostaining approaches with brightfield, epifluorescent, or confocal imaging of serial tissue sections (40–42) or whole-mount tissues combined with tissue clearing strategies (26, 43–45) provide excellent X-Y resolution and well-defined compartment labeling ability but require high reagent usage, extensive labor, and limited scalability, particularly to whole organ imaging.

To address these gaps, we have developed an accessible, high-resolution experimental-computational pipeline for anatomically accurate 3D reconstruction of relevant molecular transport barriers of murine lymph nodes (**Fig. 2**). This method integrates optimized vibratome sectioning, multiplexed immunofluorescence labeling, confocal imaging, and custom MATLAB-based automated alignment and segmentation algorithms. Our approach not only achieves precise, compartment-specific reconstructions of LN lobules and vasculature but also significantly reduces reagent use and manual labor, making it scalable and resource-efficient. Validated against manually segmented ground truth, our automated pipeline demonstrates high accuracy, ensuring robust 3D models suitable for downstream computational applications, such as spatial pharmacokinetic modeling and predictive simulations of drug distribution. Ultimately, this hybrid methodology lays the foundation for a range of computational-based methods, including digital twin-based anatomically informed modeling and therapeutic design, applicable across a broad spectrum of biological tissues and disease contexts.

**Fig. 2.**
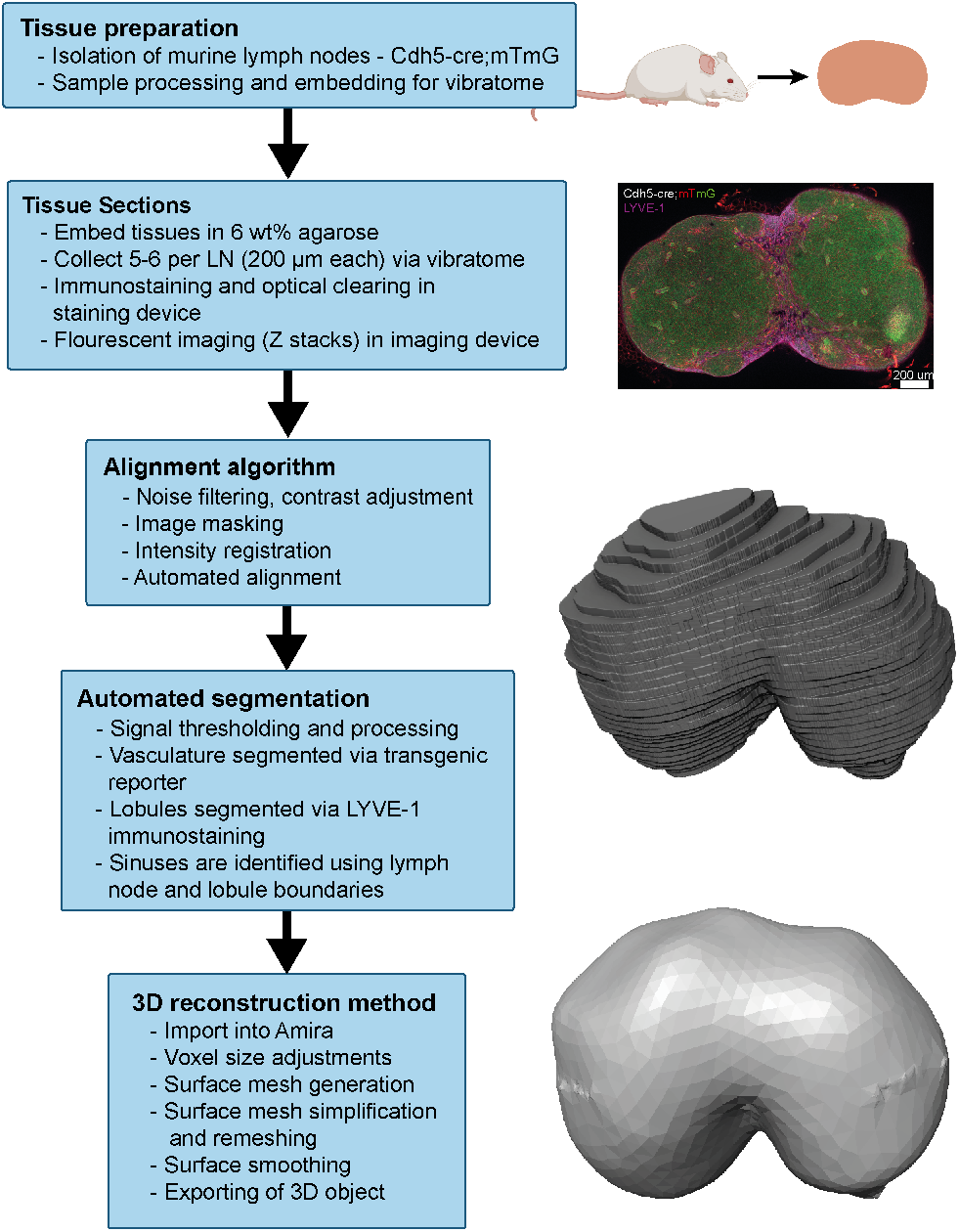
Custom pipeline to 3D reconstruct murine lymph node compartments of interest. Overview of steps on the left with visualization of output on the right.

## Materials and Methods

### Animal model and sample processing

Transgenic mice (Cdh5-Cre; mTmG, Jax Laboratories, #017968, #007676) were used for the isolation of lymph nodes (**Fig. 3A**). These mice express GFP in all VE-cadherin expressing cells, such as vasculature and lymphatic endothelial cells (LECs), and RFP in all other cells. The inguinal, popliteal, and brachial lymph nodes were excised from adult mice with wide margins, including surrounding adipose tissue. The lymph nodes were further dissected under a dissection microscope to carefully remove fat and fascia from the lymph node tissue (**3BC**). Lymph nodes were fixed overnight (approximately 18 hours) at 4ºC in 4% paraformaldehyde (PFA) with 0.1% Triton-X (Sigma-Aldrich, St. Louis, MO). After washing in 1X phosphate buffer saline (PBS), samples were transferred to 10×10×5 mm vinyl cryo-molds (Tissue-Tek, Sakura, Torrance, CA). and a 6 wt% low-melt agarose (IBI Scientific, Dubuque, IA) solution was added to the sample molds and allowed to solidify on ice to embed the lymph nodes in agarose for sectioning.

**Fig. 3.**
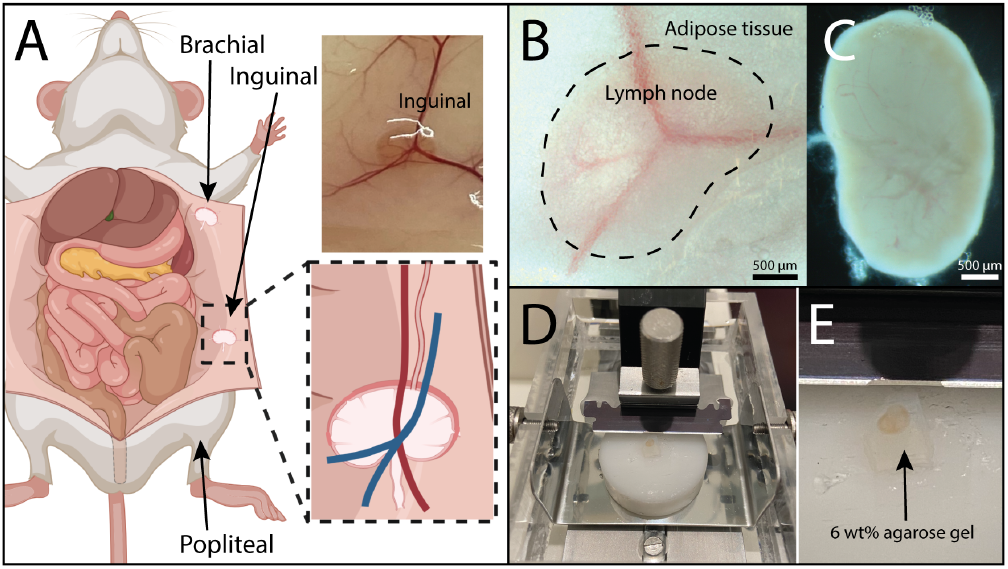
Lymph node dissection and sectioning. **A)** Brachial, inguinal, and popliteal lymph nodes are collected from the subcutaneous fat layer of mice. **B)** Isolated tissues are **C)** gently removed from surrounding adipose tissue to expose the capsule. Lymph nodes were embedded in agarose and **D**,**E)** sectioned via vibratome

### Sectioning and collection of serial slices

Gelled agarose samples were removed from the molds, and trimmed to leave approximately 2-3 mm margins from the tissue (**Fig. 3E**). A corner of the agar block was removed (**Fig. 3E**) for identification of proper sample orientation. The samples were sectioned using a vibratome (Campden 752M Vibroslice) with a section thickness of 100 or 200 *µ*m, bath advance of 0.24 mm/s, and a blade frequency of 1800 RPM (30). All slices were collected and stored in PBS in sequential order in a 48-well plate.

### Staining and optical clearing

To identify sinus boundaries, anti-LYVE1 antibody (Novus Biologicals, Centennial, CO) was used to label lymphatic endothelium. Custom staining devices were produced to efficiently immunofluorescently stain the tissues using small volumes of antibody solution, as previously described(46). First, 3D printed negative molds were generated, with 5 section slots per device(**Fig. 4A**). The fabricated pieces consisted of a 3 mm central channel to allow antibody solution to coat the tissue on either side, and five 3 mm slots on either side to hold the agar surrounding the tissue in place. Multiple negative molds were placed sequentially according to number of slots required, such as 3 molds for 15 total vibratome sections. Polydimethylsiloxane (PDMS) at a base:curing agent ratio of 20:1 was poured over the negative molds, degassed, and baked overnight at 60°C. PDMS structures were removed from the mold, and plasma-bonded to glass coverslips to produce watertight wells (**Fig. 4B**). Sections were placed orthogonally into the well slots in sequential order to ensure antibody access to both sides of the tissue (**Fig. 4C**). Samples were incubated on a rocker at 4 °C for 48 hours with the primary antibody, followed by 48 hours of incubation with the secondary antibody in the same well. After antibody labeling, tissues were incubated at room temperature with optical clearing agent (FocusClear, CelExplorer Labs, Hsinchu, Taiwan) for 2 hours (**Fig. 4DE**).

**Fig. 4.**
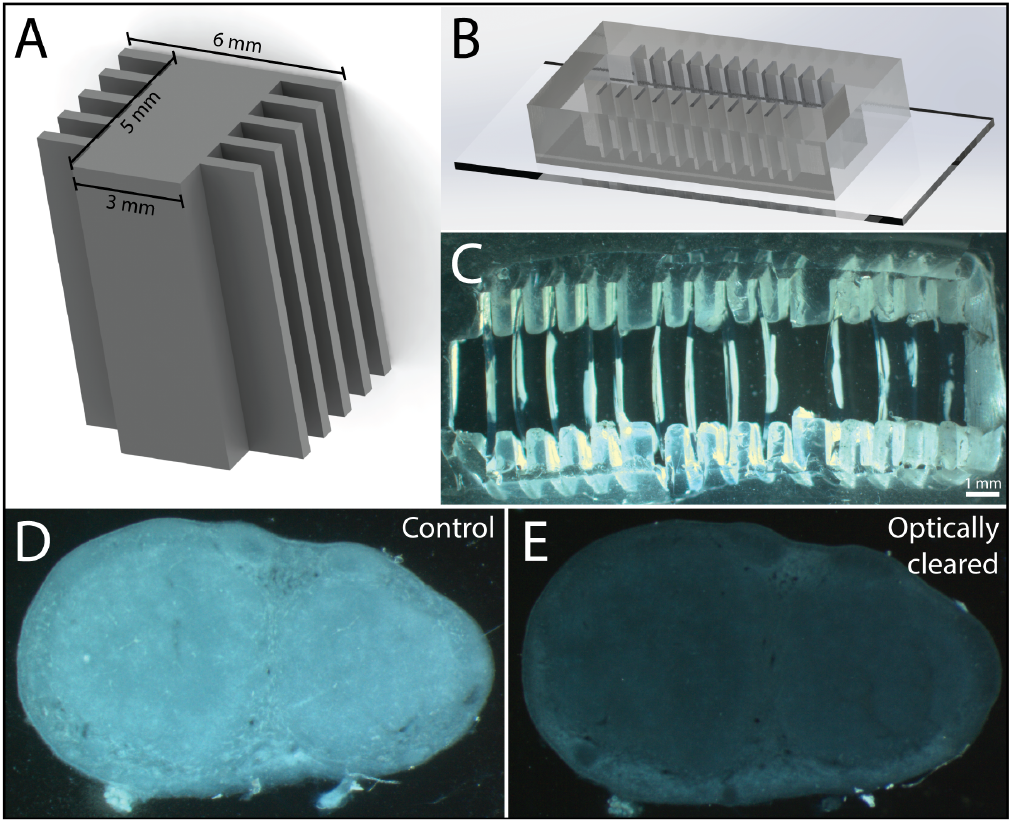
Serial sections are immunostained and optically cleared to enable high resolution z-stack imaging. **A)** 3D printed molds were used to make **B)** tissue staining devices from PDMS. **C)** Lymph node sections were placed orthogonally into the wells using the slots in the device and stained with antibody solutions. Tissues before and **E)** after optical clearing using FocusClear.

For proper placement of thick tissues on the microscope, custom glass/PDMS imaging cassettes (Fig. **5A**) were used. A 100 mm petri dish was spin-coated with PDMS to a height of 200 *µ*m (5 grams 1:20 PDMS, 300 *RPM* for 30 s, Spin Coater WS-650MZ-23NPPB, Laurell Technologies, Lansdale, PA). After curing overnight, this thin coat of PDMS was cut into a large rectangle with approximately four 4×5 mm slots. After optical clearing, the samples were transferred to the custom imaging cassettes (Fig. **5BC**), and another glass slide was placed on top of the wells to keep the sections flat against the bottom glass and prevent evaporation of the clearing solution (**Fig. 5C**). Sections were imaged using a confocal microscope (LSM800, Zeiss, EC PlnN 5x/0.16 Ph1 DIC0, Pln Apo 10x/0.45 Ph1 DICII and LD PlnN 20x/0.4 Ph2 DICII). Lasers lines of 488, 561, and 640 nm were used for imaging.

**Fig. 5.**
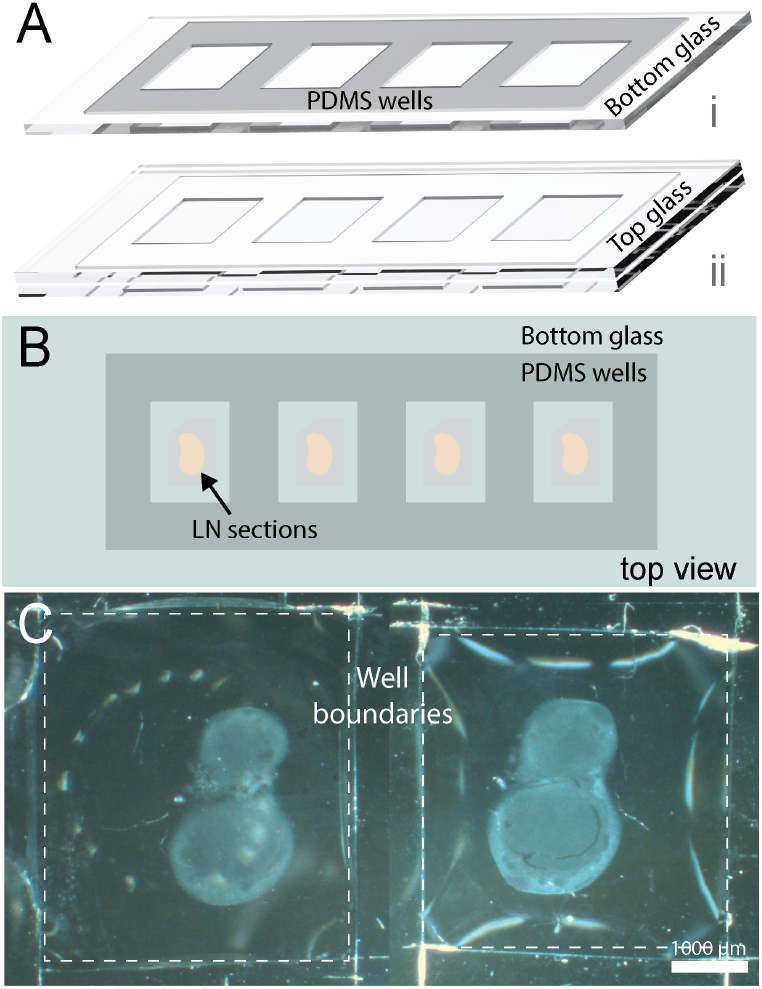
Labeled tissue sections are imaged using a custom imaging cassette. **A)**3D rendering of imaging cassette. Imaging cassettes are produced by plasma bonding a thin 200 *µ*m PDMS to a glass coverslip and then enclosing the with another glass coverslip. **B)** Schematic and **C)** representative image of tissue section in imaging device.

### Development of the alignment code

Fluorescent image stacks of each serial section were collected to identify HEVs and sinus boundaries and aid in segmentation (**Fig. 6A**). Serially imaged sections result in minor translational and rotational variations. Due to the misaligned orientation, an alignment algorithm was developed to eliminate any positional discrepancies. MATLAB (R2020B) code was developed to align the stack of lymph node images based on regions of similar intensity within the tissue (**Fig. 7**). The output was an aligned set of z-stacks for an entire lymph node. The entire sequence of TIF files was loaded into the workspace. RGB images were read from the files in pairs, starting from the first two files in the TIF sequence. The RGB images were converted to grayscale, noise-filtered, and then contrast-adjusted. For each pair of images, a freehand tracing tool was used to manually trace around the boundaries of the lymph, thereby masking out artifacts. Utilizing MATLAB’s intensity-based registration function, the pair of masked grayscale images was aligned. The last Z location in each physical tissue section was aligned to the first Z location in the subsequent stack. This transformation matrix was applied to the entire original stack for each individual channel.

**Fig. 6.**
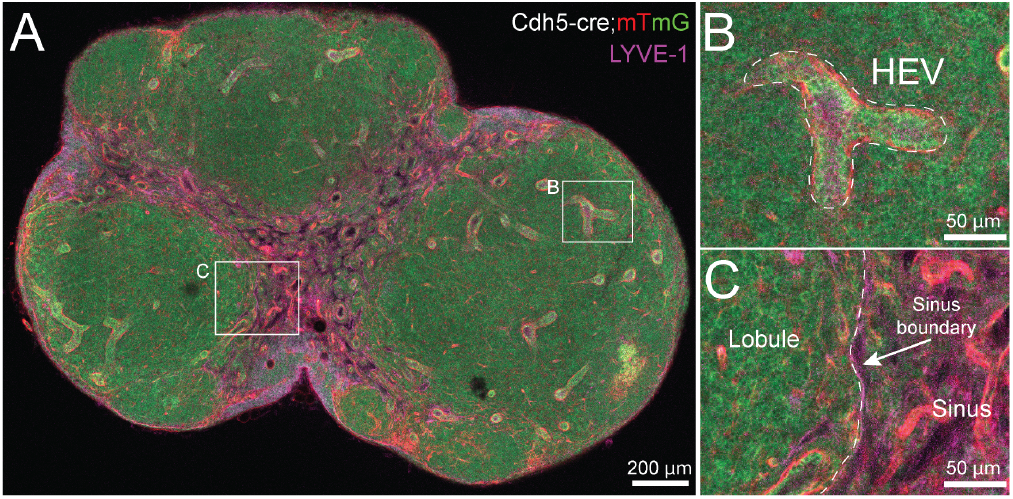
High resolution image stacks are acquired of each section. **A)** Cross section of a Cdh5-cre;mTmG mouse lymph node counterstained with anti-LYVE-1 antibody. Using this labeling scheme, both **B)** HEVs and **C)** lobule-sinus boundaries can be identified and segmented.

**Fig. 7.**
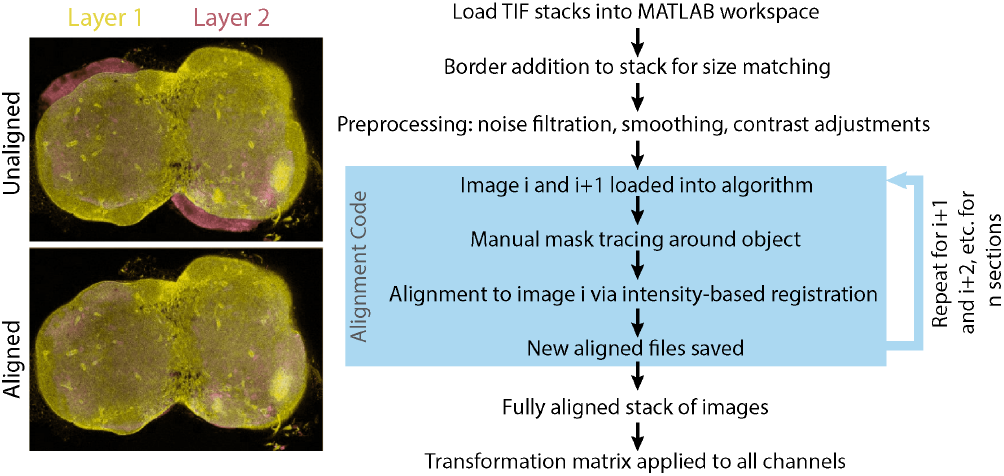
Serially acquired lymph node images aligned (left) using a custom MATLAB alignment algorithm (right).

### Image segmentation

#### Manual image segmentation

The two volumes of interest in the LN, the lobule and the HEV compartment, were manually segmented using Amira (Thermo Fisher Scientific, Waltham, MA). The aligned image stack was imported into an Amira project window, and the voxel sizes were corrected based on the pixel size-to-length conversion. A median filter was applied to remove extraneous noise from the image stack, and thresholding was used to identify the VE-cadherin signal. The paintbrush tool was used to hand-segment the vasculature and the lobule. The lobule was identified by tracing the boundary of the sinuses from the cortex to the medulla. The vasculature was identified by tracing the punctate ring structures in the lobule regions. For both tissue compartments, islands (objects smaller than 50 pixels^2^) were removed, holes (voids smaller than 50 pixels^2^) were filled, and the boundaries were processed via a smoothing function.

#### Development of an automated segmentation pipeline

A novel computational pipeline was designed using conventional computer vision techniques in MATLAB to segment the aligned z-stacks (**Fig. 8**). Due to the uncommon recombination of the CDH5-cre mouse and the progenitor pool, a specialized automated segmentation pipeline was developed specifically for this sample type.

**Fig. 8.**
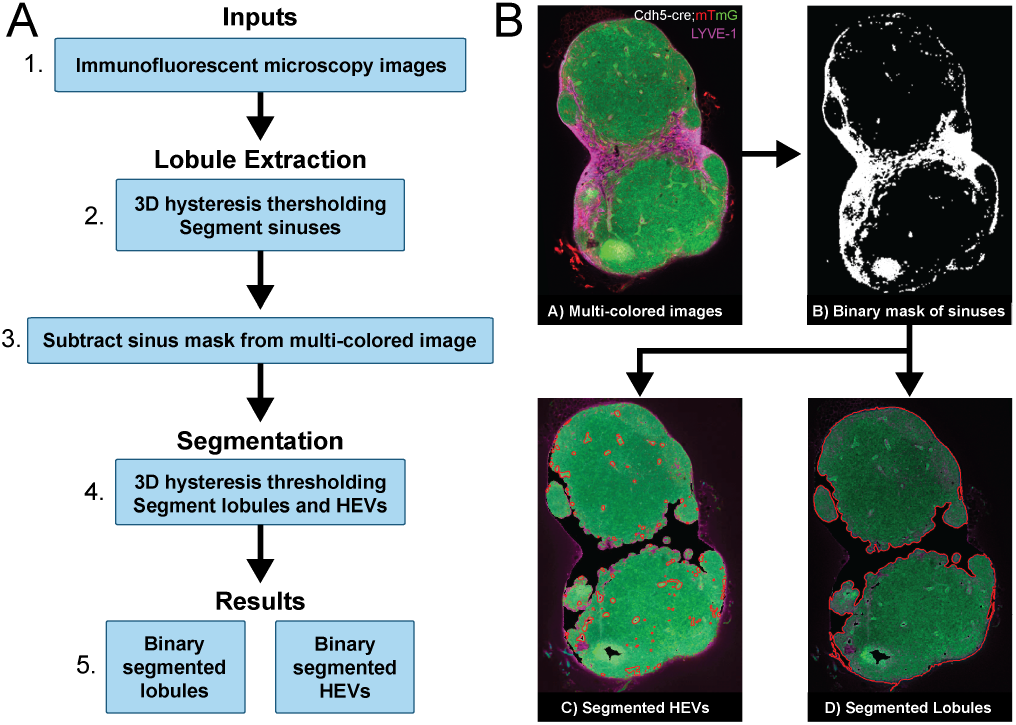
Vascular and lobule compartments are isolated throughout the entirety of the tissue stack using a custom automated segmentation pipeline via MATLAB. **A)** Flowchart of autosegmentation pipeline. **B)** Visual representation of images at different steps of the pipeline.

#### Segmenting the HEV vasculature via the automated pipeline

Images from all three channels, as well as the merged image, were imported into MATLAB in .TIF formats (**Fig. 6A**). A hysteresis thresholding algorithm (47) was applied to the LYVE1 channel image to segment the brightest of its signals, which represent the subcapsular sinuses of the lymph node (**Fig. 6A2**). Dilation and erosion functions were applied to the segmented sinuses to create a smooth and fully connected mask. This mask was then subtracted from the original multicolor image (**Fig. 6A3**). The resulting image comprised three segmented regions: a) the lymph node lobule, b) HEVs, and c) background.

For vasculature segmentation, images had contrast adjustments, median filtering, and a 2D order statistic filter was applied to aid in distinguishing the HEVs. A second round of 3D hysteresis thresholding was subsequently applied, using adjusted thresholds, to segment the HEVs based on their amplified intensities (**Fig. 6A4**). Using MATLAB’s built-in functions, the objects with the most relevant areas were extracted and identified as HEVs (**Fig. 6A5**).

These objects were then scored using a brute force algorithm to evaluate the proximity of objects in adjacent slices, thereby differentiating genuine HEVs from artifacts. The underlying rationale was that if a segmented object represents an HEV, it is expected to exhibit elongation across neighboring slices, unless it is an in-plane HEV.

#### Segmenting the lobules via the automated pipeline

The original images were processed using the same procedure outlined in the previous section through subtraction of subcapsular sinuses (**Fig. 6A3**). A median filtering algorithm was applied to remove abnormalities as a preparation step for segmentation. Subsequently, the 3D hysteresis thresholding was employed to obtain distinct outlines of the LN lobules (**Fig. 6A4**). This pipeline was then applied to the entire stack of images. Both autosegmented regions were compared to hand-segmented ground truths for each sample to confirm proper labeling.

### 3D reconstruction

The output from the automatic or manual HEV or lobule segmentation was imported into Amira, with voxel size adjustments made according to the relevant experimental data. The stacks were corrected for autosegmentation artifacts using the segmentation tool and subsequently reconstructed using Amira’s surface generation tools. The resulting reconstruction was saved as an STL file.

## Results

### Manual segmentation establishes ground truth for validation

To establish ground truth for validation, we manually segmented lobules, HEVs, and lymphatic sinuses from aligned image stacks using Amira. These compartments are critical for drug transport modeling yet are rarely co-localized in existing 3D lymph node reconstructions. For the capsule, the entire tissue was segmented using signal intensity thresholding. This was made possible by the clean tissue resection process. Any extraneous signal from surrounding adipose tissue was removed with the hand brush tool. For hand segmentation, we used lymph nodes from Cdh5-Cre x mTmG mice immunostained with LYVE1. Vasculature segmentation was based on strong GFP signal and confirmed with td-Tomato signal, as many of the perivascular cells and reticular fibroblasts created a boundary around the vasculature. For the lobules, LYVE1 immunostaining provided clear boundaries for the sinuses, allowing for the identification of lobule boundaries from the cortex to the medulla. In applications for mapping drug transport in the lymph node, the specific geometries of the sinuses are less relevant as the lymph is assumed to be a well-mixed fluid. However, if desired, the sinuses can be segmented straightforwardly by subtracting the lobule segmentation from the capsule segmentation. While sinus geometry is less critical for well-mixed lymph compartment models, this structure may be important for modeling inflamed or diseased states.

### Automated segmentation achieves high accuracy for both compartments

#### Lobule

To evaluate the level of accuracy of the automated segmentation pipeline’s outputs with respect to the hand-segmented ground truth, we used multiple types of image similarity metrics to match the specific goals of segmenting different compartments. As the lobules are studied as isolated 3D objects, the accuracy of the volumetric space is most important in their reconstruction, therefore, we chose to evaluate the similarity of the lobule segmented versions via the Sørensen-Dice (SD) index (48, 49). The SD index measures the spatial overlap between two nominal segmented images, A and B, by concentrating on their respective target regions as follows:

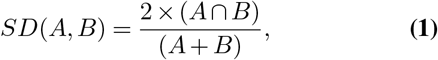

The calculations were performed in MATLAB with the two segmented versions as binary mask image inputs. The SD score result ranges from 0 (no overlap) to 1 (fully matched) and is reported as a decimal. **Fig. 9** illustrates the calculation of the SD score for four representative slices. Across all samples, mean Sørensen-Dice indices exceeded 0.85 (range:0.86-0.94), indicating high spatial overlap and validating the automated pipeline for lobule segmentation.

**Fig. 9.**
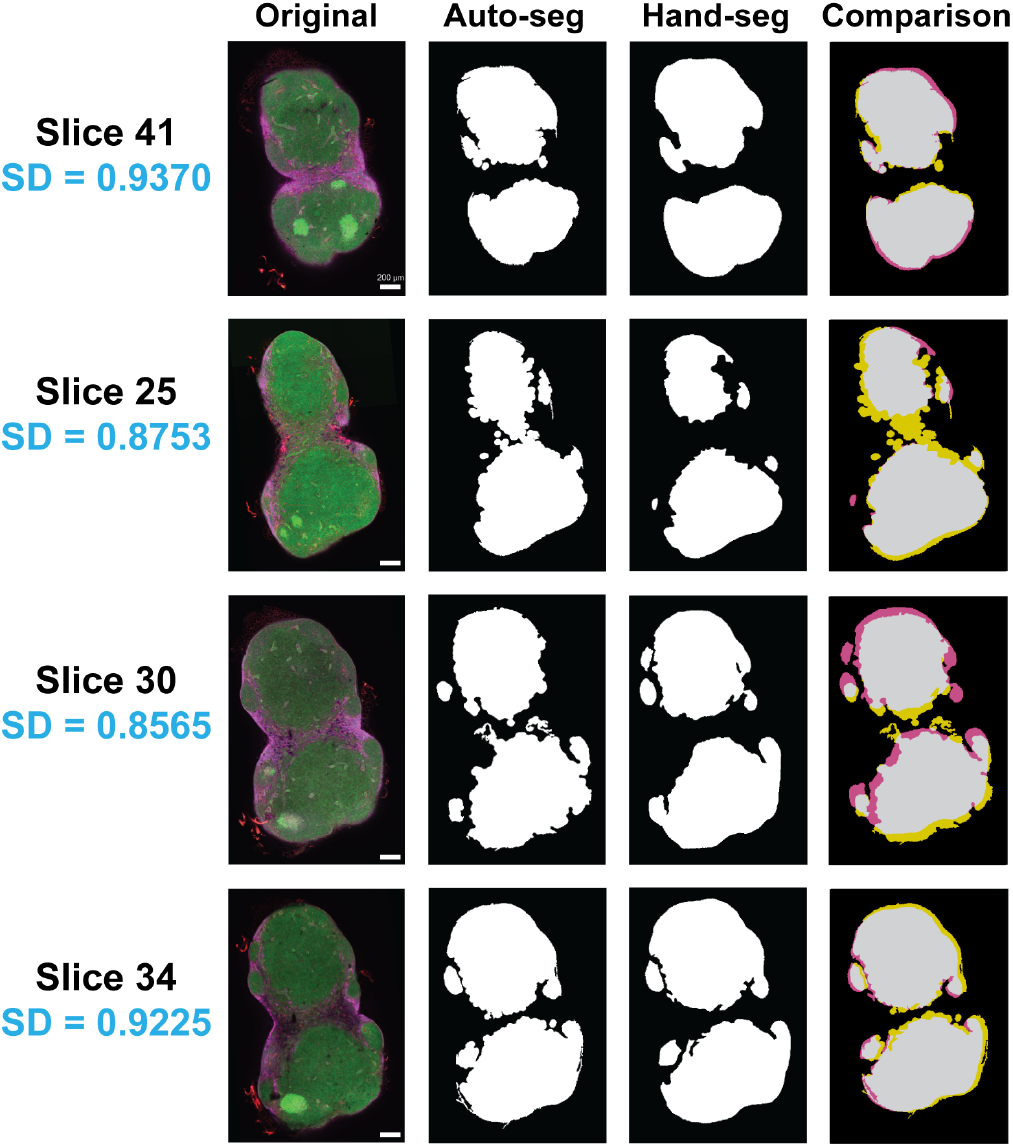
Autosegmentation vs. hand-segmentation of the lobule. Visual representation of results from randomly selected slices. The Sørensen-Dice (SD) index compares the accuracy of the automated segmentation algorithm relative to the ground truth (hand-segmented by a human expert). In the last column (Comparison) purple represents the negative false area, yellow represents the positive false area, and gray represents the true area.

#### HEVs

As HEVs have a significant tortuosity relative to lobules, and for drug transport models they act as sources for drug flux from the systemic circulation into the lymph node lobule, we chose to maximize the segmentation accuracy for surface area as opposed to volumetric accuracy. To do so we compared the outline contours of the segmented HEVs from our pipeline relative to hand-segmented ground truths based on a contour-based matching score (50):

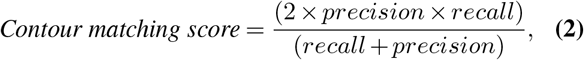

where “precision” is defined as the ratio of true positive points along the segmentation’s boundary (ground truth) to the total points along the predicted boundary, and “recall” is the fraction of true positive points along the predicted segmentation’s boundary from the total points along the ground truth boundary. As such, this metric quantifies the ability to detect all relevant points along the ground-truth boundary. The overall score is calculated by taking the harmonic mean of the precision and recall values. This metric evaluates the automated segmentation’s correspondence with the hand-segmentation (ground truth), while accounting for a distance error tolerance.

The calculations were done in MATLAB with the two segmented versions of the HEV vasculature as binary mask image inputs. The comparison of the same four representative image slices for HEV vasculature segmentation using the contour-matching score (**Fig. 10**) shows up to 77% matching with the ground truth. This demonstrates the fidelity of our automated algorithm is more than 70% with respect to what a human expert can determine as the HEV vasculature, which is lower than the corresponding score for lobules, but adequate for our further purposes of drug delivery simulations.

**Fig. 10.**
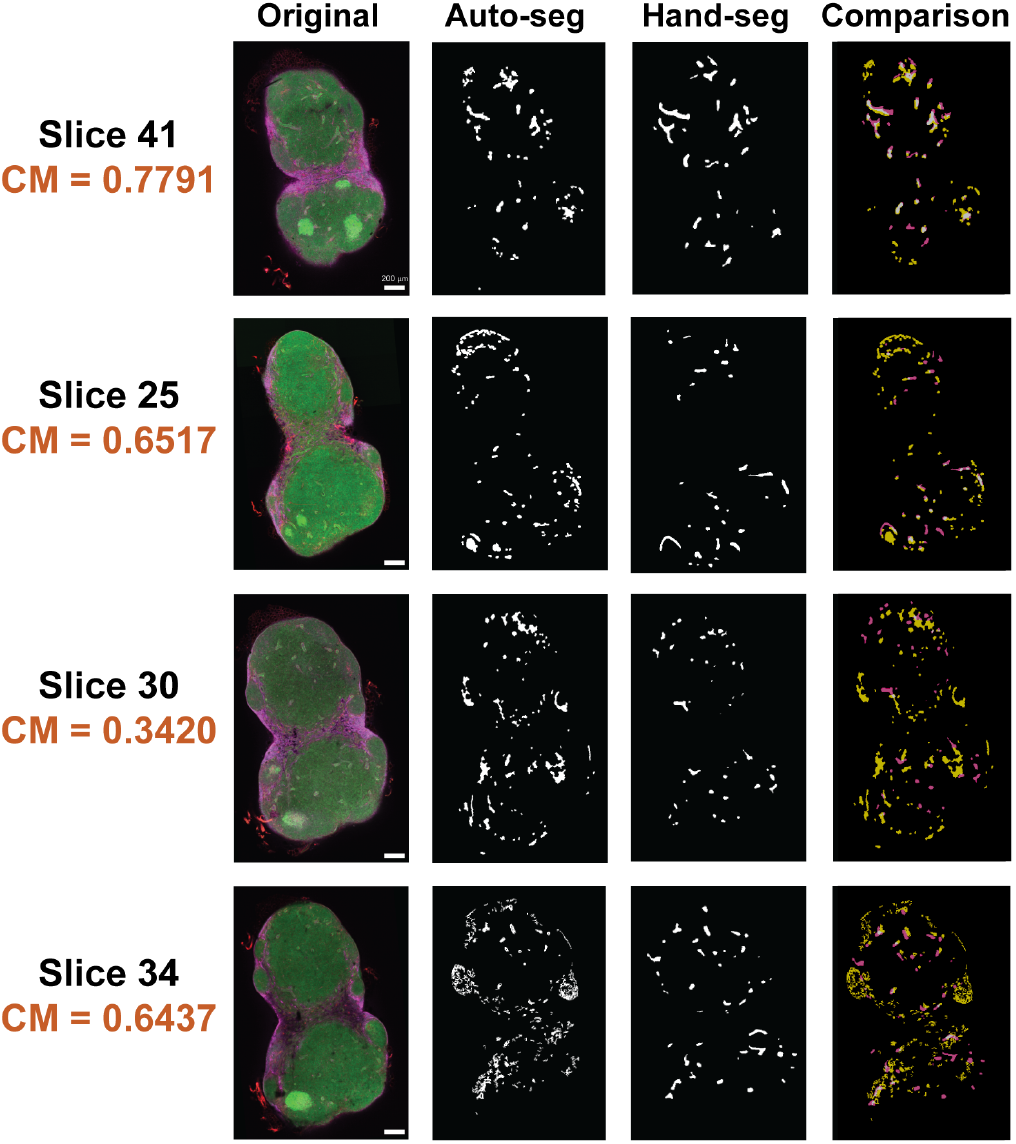
Autosegmentation vs. hand-segmentation of the HEVs. Visual representation of results from randomly selected slices. A contour matching (CM) metric is reported for the relative congruence between the autosegmentation output and the hand-segmented final result. In the last column (Comparison) purple represents the negative false area and yellow represents the positive false area.

### Reconstructed 3D geometries provide anatomical domains for geometric analysis and computational modeling

Once the vasculature and lobules were segmented, Amira was used for the 3D-reconstruction of the lymph node (**Fig. 11**). Segmented Tiff stacks were uploaded to Amira, and the appropriate Z voxel size was incorporated to represent the confocal step size. Whereas hand segmentations produced largely clean boundaries, a median filter with a pixel size of 3 was applied to smooth any minor human errors. A surface was generated for the lobule and vasculature. Through validation of different step sizes, 10 *µ*m steps between serial images allowed for optimal surface generation in Amira (**Fig. 11A**). This resolution in Z was found to sufficiently reduce information loss through the tissue for accurate interpolation. For visualization, the surface of the lobule was remeshed with 1% of the original surface triangles, followed by use of a surface smoothing function to remove imaging and alignment artifacts. The volume overlap between a stack of segmented images, with a section width of 10 *µ*m, and the generated surface was measured to be greater than 97% for all samples, resulting in a ∼3% overestimation of the segmented regions. Additional studies indicated that the overestimation decreases with smaller Z step sizes; however, computational and experimental time and expense increase significantly.

**Fig. 11.**
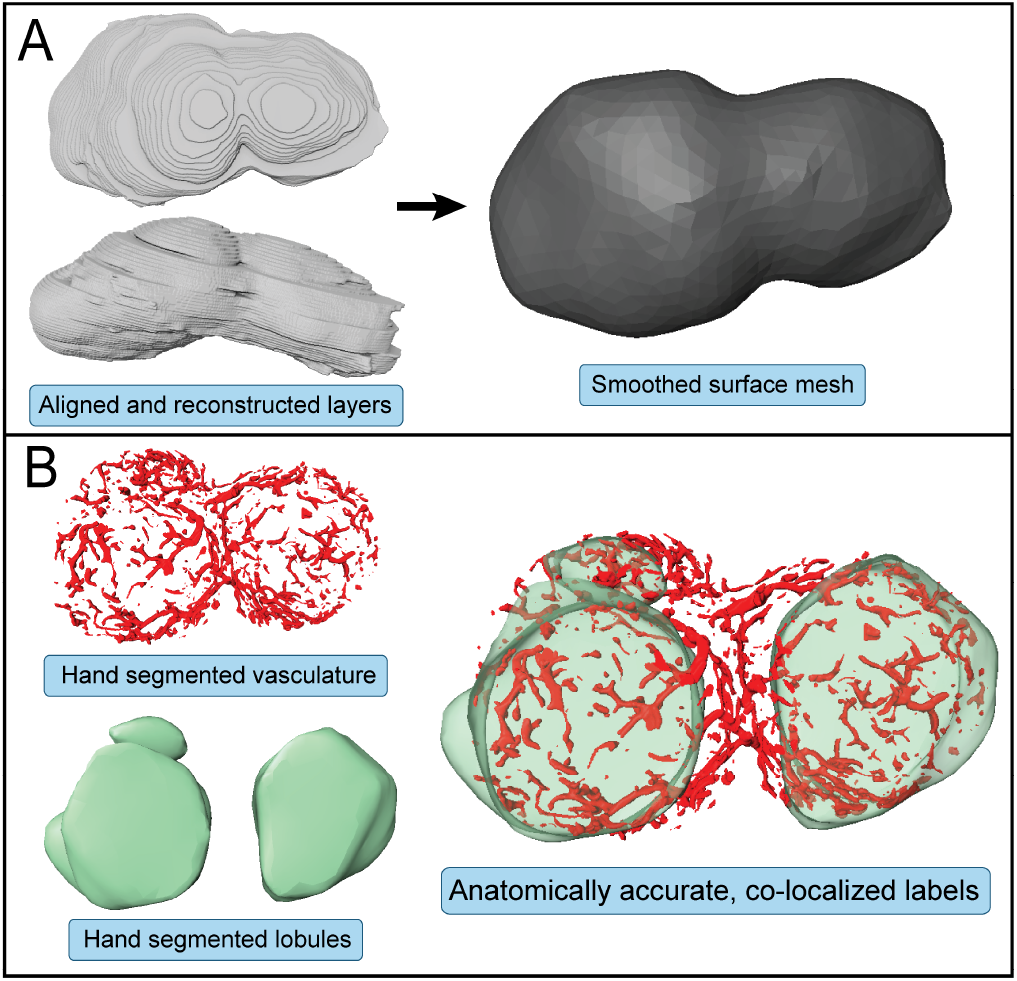
Segmented lymph node compartments were 3D reconstructed. **A)** After serial tissue sections were segmented and exported, Amira was used to create a 3D volume and generate a surface mesh. **B)** The two compartments of interest, HEVs and lobules, were reconstructed as 3D objects. These objects and their geometric parameters can then be used to be able to study interactions between the lobule parenchyma and both fluid compartments.

The resulting 3D reconstructed domains can be processed as separate 3D objects or combined using the global coordinate systems shared with each domain (**Fig. 11B**). The resulting 3D reconstructed lymph nodes match what is seen from empirically collected immunofluorescence images and are consistent with the literature (16). These geometric parameters can be directly used as inputs for finite element modeling of drug distribution. For a small number of lymph nodes, this protocol is efficient at 3D anatomical reconstruction for analysis; however, automated sectioning and imaging systems are required for a higher throughput.

## Discussion

This work addresses a critical bottleneck in computational modeling of lymph node drug transport; the lack of accessible methods for generating anatomically accurate 3D tissue reconstructions with standard laboratory microscopes. Our validated pipeline reduces reagent costs by ∼88% and manual labor by ∼97% compared to traditional serial sectioning **(Table 1)** while achieving expert-level segmentation accuracy (Sørensen-Dice >0.93 for lobules, contour-matching scores up to 77% for HEVs). This efficiency gain transforms anatomical reconstruction from a months-long endeavor requiring specialized expertise into a routine technique that can be performed in 1-2 days using standard laboratory equipment. By demonstrating that high-fidelity reconstructions can be generated without cleanroom facilities, specialized tissue clearing protocols, or vacuum sealing systems, we lower barriers to entry for laboratories across disciplines, enabling population-level studies that were previously infeasible.

**Table 1.** Comparison of LN preparation, imaging, and segmentation methods.

| Parameter | Our Method | Traditional Manual |
| --- | --- | --- |
| **Sample Preparation** |  |  |
| Embedding + Sectioning | 1 hr | 4 hr |
| Number of sections | ~15 | ~150 |
| **Staining and Mounting** |  |  |
| Setup time | 30 min | — |
| Staining time | 1.5 hr | 5 hr |
| Time per slice | 6 min | 2 min |
| Mounting slides | — | 0.5 min/slice |
| **Imaging** |  |  |
| Setup time | 30 min | — |
| Time per section | 15 min | 4 min |
| **Segmentation** |  |  |
| per section | <1 min | 1.25 hr |
| **Total per Section** | 30 min | 1.38 hr |
| **Total per Lymph Node** |  |  |
| Full Prep/Imaging/Segmentation | **7.5 hr** | **207.75 hr** |
| Reagent Cost | **~\$45** | **~\$370** |

Compared to traditional imaging methods such as microcomputed tomography (µCT), magnetic resonance imaging (MRI), whole-mount optical clearing, or conventional serial histological sectioning, our pipeline offers distinct advantages in balancing resolution, compartment-specific labeling, and accessibility. Unlike µCT and MRI, which lack multiplexed labeling or whole-mount clearing (51, 52), which is limited to small tissues, our approach combines compartment-specific immunolabeling with scalable throughput suitable for larger lymph nodes relevant to disease modeling. Most critically, the method requires only standard equipment: a vibratome, a confocal microscope, and MATLAB software. Custom staining devices reduce antibody consumption by >90%, decreasing per-sample costs from ∼$370 to ∼$45 **(Table 1)**. Our automated alignment and segmentation algorithms, while developed for lymph nodes, employ generalizable computer vision techniques (intensity-based registration, hysteresis thresholding, morphological operations) adaptable to other compartmentalized tissues including spleen, gut-associated lymphoid tissues, or vascularized tissues and tumors (53–55).

The anatomically accurate digital geometries generated by this pipeline directly enable several research applications without requiring additional experimental measurements (56–58). Finite element modeling using these geometries could predict which drug formulations will achieve therapeutic concentrations throughout heterogeneous tissue architecture, identifying combinations most likely to suppress viral reservoirs in HIV or penetrate metastatic niches. Population-level studies reconstructing lymph nodes from multiple individuals could determine whether anatomical variability (lobule volume, HEV network density, sinus architecture) correlates with observed differences in therapeutic response,potentially identifying anatomical biomarkers predictive of treatment efficacy. Serial reconstructions from disease models could quantify how pathological remodeling alters tissue architecture, enabling time-dependent simulations that predict how structural changes affect drug penetration as disease progresses. Understanding the spatial distribution patterns of immunotherapies based on anatomical features could inform dosing strategies and identify architectural barriers that limit efficacy. These applications leverage the reconstructed geometries as computational input domains, demonstrating the immediate translational utility of our pipeline for spatial pharmacokinetic modeling and therapeutic optimization.

The choice of validation metrics reflects intended applications wherby volumetric accuracy is critical for lobules, which define fluid compartments in transport models, while surface accuracy is paramount for HEVs, which govern blood-to-tissue exchange rates. The generated 3D models provide the spatial precision required for mechanistic simulations, addressing significant gaps in existing computational approaches that rely on idealized geometries (e.g., spheres, cylinders, homogeneous domains) and fail to replicate experimentally observed spatial drug gradients. This is particularly relevant in lymph node-resident diseases such as HIV and metastatic cancer, where therapeutic exclusion is mediated by tissue-specific, dyanmic barriers like HEVs and sinus-parenchyma interfaces absent from simplified models (59). Our reconstructions enable the generation of personalized or population-based anatomical models that can inform more accurate spatial pharmacokinetic simulations, providing mechanistic explanations for therapeutic failures and guiding drug delivery optimization (29).

Despite these advancements, there are opportunities for future development. We demonstrated reconstruction of two critical compartments (lobules and HEVs); additional structures, including B-cell follicles, T-cell zones, and medullary cords, can be incorporated by adding compartment-specific markers without fundamental pipeline changes. The current workflow requires a full day of labor and 2-3 days of incubation time per lymph node. Most of this effort is focused on LN preparation, staining, and imaging. Further automation of sectioning and imaging could reduce processing time to <24 hours, enabling larger population studies. Future iterations of our pipeline will incorporate more specific cellular immunostaining markers, enhanced segmentation algorithms including active contour models, and extend applicability to larger samples such as human and non-human primate tissues. The outputs of this pipeline are 3D object files that are compatible with all standard meshing software packages. By importing these meshes of lymph node vasculature and lobules into a finite volume method (FVM) model of the lymph node, we hope to computationally validate key drug, cell, and pathogen transport parameters against experimental data (60). An accurate spatial model of drug transport in the lymph node will allow us to develop a valuable predictive tool for drug transport in lymph nodes of varying geometries and disease states across many drug classes.

By making anatomical reconstruction accessible through standard equipment and open-source code, we provide a blueprint for laboratories across disciplines to integrate spatial modeling into their research programs. The modular design enables adaptation to other tissues and species, positioning this pipeline as a foundation for creating digital anatomical models across diverse biological and biomedical applications (61–64). We anticipate that this accessible platform will enable population-level anatomical studies correlating tissue architecture with therapeutic outcomes, research that was previously infeasible due to throughput and cost limitations. As computational modeling becomes increasingly integrated into biomedical research, accessible methods for generating anatomically accurate input geometries will be essential tools for the research community.

## Conclusion and Future Directions

We developed a robust, scalable, and highly accurate pipeline for anatomical 3D reconstruction of murine lymph nodes that achieves expert-level segmentation accuracy (Sørensen-Dice >0.93 for lobules, contour-matching up to 77% for HEVs) while reducing reagent costs by ∼88% and manual labor by ∼97% compared to traditional approaches. By integrating optimized vibratome sectioning, multiplexed immunofluorescence, automated segmentation, and rigorous validation, this method provides anatomically accurate digital geometries suitable for computational modeling of drug transport and immune cell dynamics using only standard laboratory equipment. The approach is adaptable to other tissues and species, with future extensions to human and non-human primate lymphoid specimens, additional compartments (B-cell follicles, T-cell zones, conduits), and integration with computational simulations to validate predictive models. Ultimately, this accessible pipeline bridges experimental practices and computational modeling, enabling the emerging field of anatomically informed pharmacology, where tissuespecific geometry guides therapeutic design for lymph node-resident diseases, including HIV and metastatic cancer.

## ACKNOWLEDGEMENTS

The authors thank Katherine M. Nelson, Ph.D., for reviewing and commenting on the manuscript. This work was supported in part by grants from the National Institutes of Health (R21AI157889-JPG, RZ; T32GM142603-YM), the National Science Foundation (CBET 1943686-JPG), and the State of Delaware through the Delaware Biotechnology Institute’s Bioscience CAT (JPG). The BioRxiv template was adapted from the Henriques lab.

## COMPETING FINANCIAL INTERESTS

There are no conflicts of interest.

## DATA AND CODE AVAILABILITY

MATLAB code is available at https://github.com/Gleghorn-Lab/LymphNodeReconstruction

## AUTHOR CONTRIBUTIONS

Conceptualization (MJD, YM, RZ, JPG), Methodology (MJD, YM, ATK), Investigation (MJD, YM, ATK), Validation (MJD, YM, RZ, JPG), Visualization (MJD, YM, ATK), Formal Analysis (MJD, YM, RZ, JPG), Writing – Original Draft (MJD, YM), Writing – Review & Editing (MJD, YM, ATK, RZ, JPG), Supervision (RZ, JPG), Project Administration (JPG), Funding acquisition (JPG)

## Notes

### Competing Interest Statement

The authors have declared no competing interest.

### Summary of Updates

The funding acknowledgement has been updated.

## References

1. Diana Murray, Donald Petrey, and Barry Honig. Integrating 3D structural information into systems biology. The Journal of Biological Chemistry, 296: 100562, 2021. ISSN 1083-351X. doi: 10.1016/j.jbc.2021.100562.

2. Britta A. M. Bouwman, Nicola Crosetto, and Magda Bienko. The era of 3D and spatial genomics. Trends in genetics: TIG, 38(10):1062–1075, October 2022. ISSN 0168-9525. doi: 10.1016/j.tig.2022.05.010.

3. John M. Lucocq, Terry M. Mayhew, Yannick Schwab, Anna M. Steyer, and Christian Hacker. Systems biology in 3D space–enter the morphome. Trends in Cell Biology, 25(2):59–64, February 2015. ISSN 1879-3088. doi: 10.1016/j.tcb.2014.09.008.

4. Kara E. Peak, Shelby R. Mohr-Allen, Jason P. Gleghorn, and Victor D. Varner. Focal sources of FGF-10 promote the buckling morphogenesis of the embryonic airway epithelium. 11(9):bio059436. ISSN 2046-6390. doi: 10.1242/bio.059436.

5. Laurel E. Schappell, Daniel J. Minahan, and Jason P. Gleghorn. A microfluidic system to measure neonatal lung compliance over late stage development as a functional measure of lung tissue mechanics. 142(100803). ISSN 0148-0731. doi: 10.1115/1.4047133.

6. Brielle Hayward-Piatkovskyi, Cailin R. Gonyea, Sienna C. Pyle, Krithika Lingappan, and Jason P. Gleghorn. Sex-related external factors influence pulmonary vascular angiogenesis in a sex-dependent manner. 324(1):H26–H32. ISSN 0363-6135. doi: 10.1152/ajpheart.00552.2022. Publisher: American Physiological Society.

7. Celeste M. Nelson and Jason P. Gleghorn. Sculpting organs: Mechanical regulation of tissue development. 14(1):129–154. ISSN 1523-9829, 1545-4274. doi: 10.1146/annurev-bioeng-071811-150043.

8. Katherine M. Nelson, Daniel J. Minahan, Vonetta L. Edwards, Ian J. Glomski, David J. Delgado Diaz, Keena Thomas, Forrest C. Walker, Patrik M. Bavoil, Isabelle Derré, Alison K. Criss, Jacques Ravel, and Jason P. Gleghorn. A microphysiologic model of the cervical epithelium recapitulates microbial, immunologic, and pathogenic properties of sexually transmitted infections. ISSN: 2692-8205 Pages: 2025.07.21.665989 Section: New Results.

9. Daniel J. Minahan, Katherine M. Nelson, Filipa Ribeiro, Bryan J. Ferrick, Alexandra M. Zurzolo, Kira Byers, and Jason P. Gleghorn. Democratizing organ-on-chip technologies with a modular, reusable, and perfusion-ready microphysiological system. Pages: 2025.04.30.651503 Section: New Results.

10. Joshua T. Morgan, Wade G. Stewart, Robert A. McKee, and Jason P. Gleghorn. The mechanosensitive ion channel TRPV4 is a regulator of lung development and pulmonary vasculature stabilization. 11(5):309–320,. ISSN 1865-5033. doi: 10.1007/s12195-018-0538-7.

11. Davide Ambrosi, Martine Ben Amar, Christian J. Cyron, Antonio DeSimone, Alain Goriely, Jay D. Humphrey, and Ellen Kuhl. Growth and remodelling of living tissues: perspectives, challenges and opportunities. Journal of the Royal Society Interface, 16(157):20190233, August 2019. ISSN 1742-5689. doi: 10.1098/rsif.2019.0233.

12. Huaiyu Shi, Chenyan Wang, and Zhen Ma. Stimuli-responsive biomaterials for cardiac tissue engineering and dynamic mechanobiology. APL Bioengineering, 5(1):011506, March 2021. ISSN 2473-2877. doi: 10.1063/5.0025378.

13. Evangelia Katsoulakis, Qi Wang, Huanmei Wu, Leili Shahriyari, Richard Fletcher, Jinwei Liu, Luke Achenie, Hongfang Liu, Pamela Jackson, Ying Xiao, Tanveer Syeda-Mahmood, Richard Tuli, and Jun Deng. Digital twins for health: a scoping review. npj Digital Medicine, 7(1):1–11, March 2024. ISSN 2398-6352. doi: 10.1038/s41746-024-01073-0. Publisher: Nature Publishing Group.

14. Jiqing Wu and Viktor H. Koelzer. Towards generative digital twins in biomedical research. Computational and Structural Biotechnology Journal, 23:3481–3488, December 2024. ISSN 2001-0370. doi: 10.1016/j.csbj.2024.09.030.

15. Xinxiu Li, Joseph Loscalzo, A. K. M. Firoj Mahmud, Dina Mansour Aly, Andrey Rzhetsky, Marinka Zitnik, and Mikael Benson. Digital twins as global learning health and disease models for preventive and personalized medicine. Genome Medicine, 17(1):11, February 2025. ISSN 1756-994X. doi: 10.1186/s13073-025-01435-7.

16. Cynthia L Willard-Mack. Normal structure, function, and histology of lymph nodes. Toxicologic pathology, 34(5):409–424, 2006. Publisher: Sage Publications.

17. Catherine E. Angel, Chun-Jen J. Chen, Oliver C. Horlacher, Sintia Winkler, Thomas John, Judy Browning, Duncan MacGregor, Jonathan Cebon, and P. Rod Dunbar. Distinctive localization of antigen-presenting cells in human lymph nodes. Blood, 113(6):1257–1267, February 2009. ISSN 1528-0020. doi: 10.1182/blood-2008-06-165266.

18. J. E. Gretz, A. O. Anderson, and S. Shaw. Cords, channels, corridors and conduits: critical architectural elements facilitating cell interactions in the lymph node cortex. Immunological Reviews, 156:11–24, April 1997. ISSN 0105-2896. doi: 10.1111/j.1600-065x.1997.tb00955.x.

19. J. E. Gretz, E. P. Kaldjian, A. O. Anderson, and S. Shaw. Sophisticated strategies for information encounter in the lymph node: the reticular network as a conduit of soluble information and a highway for cell traffic. Journal of Immunology (Baltimore, Md.: 1950), 157(2):495–499, July 1996. ISSN 0022-1767.

20. Sirpa Jalkanen and Marko Salmi. Lymphatic endothelial cells of the lymph node. Nature Reviews Immunology, 20(9):566–578, September 2020. ISSN 1474-1741. doi: 10.1038/s41577-020-0281-x. Number: 9 Publisher: Nature Publishing Group.

21. Ramon Roozendaal, Reina E. Mebius, and Georg Kraal. The conduit system of the lymph node. International Immunology, 20(12):1483–1487, December 2008. ISSN 1460-2377. doi: 10.1093/intimm/dxn110.

22. Britta Engelhardt and Hartwig Wolburg. Mini-review: Transendothelial migration of leukocytes: through the front door or around the side of the house? European Journal of Immunology, 34(11):2955–2963, November 2004. ISSN 0014-2980. doi: 10.1002/eji.200425327.

23. Friederike Pfeiffer, Varsha Kumar, Stefan Butz, Dietmar Vestweber, Beat A. Imhof, Jens V. Stein, and Britta Engelhardt. Distinct molecular composition of blood and lymphatic vascular endothelial cell junctions establishes specific functional barriers within the peripheral lymph node. European Journal of Immunology, 38(8):2142–2155, August 2008. ISSN 0014-2980. doi: 10.1002/eji.200838140.

24. A. O. Anderson and N. D. Anderson. Studies on the structure and permeability of the microvasculature in normal rat lymph nodes. The American Journal of Pathology, 80(3):387–418, September 1975. ISSN 0002-9440.

25. M C Kowala and G I Schoefl. The popliteal lymph node of the mouse: internal architecture, vascular distribution and lymphatic supply. Journal of Anatomy, 148:25–46, October 1986. ISSN 0021-8782.

26. Mohammad Jafarnejad, Matthew C. Woodruff, David C. Zawieja, Michael C. Carroll, and J. E. Moore. Modeling Lymph Flow and Fluid Exchange with Blood Vessels in Lymph Nodes. Lymphatic Research and Biology, 13(4):234–247, December 2015. ISSN 1557-8585. doi: 10.1089/lrb.2015.0028.

27. K. N. Margaris and R. A. Black. Modelling the lymphatic system: challenges and opportunities. Journal of the Royal Society, Interface, 9(69):601–612, April 2012. ISSN 1742-5662. doi: 10.1098/rsif.2011.0751.

28. Alex Schudel, David M. Francis, and Susan N. Thomas. Material design for lymph node drug delivery. Nature Reviews Materials, 4(6):415–428, June 2019. ISSN 2058-8437. doi: 10.1038/s41578-019-0110-7.

29. Noah Trac and Eun Ji Chung. Overcoming physiological barriers by nanoparticles for intravenous drug delivery to the lymph nodes. Experimental Biology and Medicine, 246(22):2358–2371, November 2021. ISSN 1535-3702. doi: 10.1177/15353702211010762.

30. Maura C. Belanger, Alexander G. Ball, Megan A. Catterton, Andrew W.L. Kinman, Parastoo Anbaei, Benjamin D. Groff, Stephanie J. Melchor, John R. Lukens, Ashley E. Ross, and Rebecca R. Pompano. Acute Lymph Node Slices Are a Functional Model System to Study Immunity Ex Vivo. ACS Pharmacology & Translational Science, 4(1):128–142, January 2021. ISSN 2575-9108. doi: 10.1021/acsptsci.0c00143.

31. Melody A. Swartz. The physiology of the lymphatic system. Advanced Drug Delivery Reviews, 50(1):3–20, August 2001. ISSN 0169-409X. doi: 10.1016/S0169-409X(01)00150-8.

32. Maria H. Ulvmar and Taija Mäkinen. Heterogeneity in the lymphatic vascular system and its origin. Cardiovascular Research, 111(4):310–321, September 2016. ISSN 1755-3245. doi: 10.1093/cvr/cvw175.

33. Jan M Orenstein, Mark Feinberg, Christian Yoder, Lewis Schrager, JoAnn M Mican, Douglas J Schwartzentruber, Richard T Davey Jr, Robert E Walker, Judith Falloon, Joseph A Kovacs, and others. Lymph node architecture preceding and following 6 months of potent antiviral therapy: follicular hyperplasia persists in parallel with p24 antigen restoration after involution and CD4 cell depletion in an AIDS patient. Aids, 13(16):2219–2229, 1999. Publisher: LWW.

34. Allen P Burke, David Anderson, Poonam Mannan, Jorge L Ribas, You-Hui Liang, John Smialek, and Renu Virmani. Systemic lymphadenopathic histology in human immunodeficiency virus-1—Seropositive drug addicts without apparent acquired immunodeficiency syndrome. Human pathology, 25(3): 248–256, 1994. Publisher: Elsevier.

35. Ryo Iwamura, Maya Sakamoto, Shiro Mori, and Tetsuya Kodama. Imaging of the Mouse Lymphatic Sinus during Early Stage Lymph Node Metastasis Using Intranodal Lymphangiography with X-ray Micro-computed Tomography. Molecular Imaging and Biology, 21(5):825–834, October 2019. ISSN 1860-2002. doi: 10.1007/s11307-018-01303-4.

36. Christopher T. Winkelmann, Said Daibes Figueroa, Tammy L. Rold, Wynn A. Volkert, and Timothy J. Hoffman. Microimaging Characterization of a B16-F10 Melanoma Metastasis Mouse Model. Molecular Imaging, 5(2): 7290.2006.00011, April 2006. ISSN 1535-3508. doi: 10.2310/7290.2006.00011. Publisher: SAGE Publications Inc.

37. Tetsuya Kodama, Shiro Mori, and Masato Nose. Tumor cell invasion from the marginal sinus into extranodal veins during early-stage lymph node metastasis can be a starting point for hematogenous metastasis. Journal of Cancer Metastasis and Treatment, 4(0):N/A–N/A, November 2018. ISSN ISSN 2454-2857 (Online)<br>ISSN 2394-4722 (Print). doi: 10.20517/2394-4722.2018.61. Publisher: OAE Publishing Inc.

38. Wolfgang Eck, Anthony I. Nicholson, Hanswalter Zentgraf, Wolfhard Semmler, and Sönke Bartling. Anti-CD4-targeted gold nanoparticles induce specific contrast enhancement of peripheral lymph nodes in X-ray computed tomography of live mice. Nano Letters, 10(7):2318–2322, July 2010. ISSN 1530-6992. doi: 10.1021/nl101019s.

39. M. Jafarnejad, A. Z. Ismail, D. Duarte, C. Vyas, A. Ghahramani, D. C. Zawieja,C. Lo Celso, G. Poologasundarampillai, and J. E. Moore. Quantification of the Whole Lymph Node Vasculature Based on Tomography of the Vessel Corrosion Casts. Scientific Reports, 9(1):13380, September 2019. ISSN 2045-2322. doi: 10.1038/s41598-019-49055-7.

40. Bin Ma, Zhuang Lin, Simon Winkelbach, Werner Lindenmaier, and Kurt E. J. Dittmar. Automatic registration of serial sections of mouse lymph node by using Image-Reg. Micron, 39(4):387–396, June 2008. ISSN 0968-4328. doi: 10.1016/j.micron.2007.03.005.

41. Arnauld Sergé, Anne-Laure Bailly, Michel Aurrand-Lions, Beat A. Imhof, and Magali Irla. For3D: Full organ reconstruction in 3D, an automatized tool for de-Donzanti, Moghadamnia et al. | 3D LN Anatomical Reconstruction ciphering the complexity of lymphoid organs. Journal of Immunological Methods, 424:32–42, September 2015. ISSN 0022-1759. doi: 10.1016/j.jim.2015.04.019.

42. Alexey Kislitsyn, Rostislav Savinkov, Mario Novkovic, Lucas Onder, and Gennady Bocharov. Computational Approach to 3D Modeling of the Lymph Node Geometry. Computation, 3(2):222–234, June 2015. ISSN 2079-3197. doi: 10.3390/computation3020222. Number: 2 Publisher: Multidisciplinary Digital Publishing Institute.

43. Qian Chai, Lucas Onder, Elke Scandella, Cristina Gil-Cruz, Christian Perez-Shibayama, Jovana Cupovic, Renzo Danuser, Tim Sparwasser, Sanjiv A. Luther, Volker Thiel, Thomas Rülicke, Jens V. Stein, Thomas Hehlgans, and Burkhard Ludewig. Maturation of lymph node fibroblastic reticular cells from myofibroblastic precursors is critical for antiviral immunity. Immunity, 38(5): 1013–1024, May 2013. ISSN 1097-4180. doi: 10.1016/j.immuni.2013.03.012.

44. Varsha Kumar, Elke Scandella, Renzo Danuser, Lucas Onder, Maximilian Nitschké, Yoshinori Fukui, Cornelia Halin, Burkhard Ludewig, and Jens V. Stein. Global lymphoid tissue remodeling during a viral infection is orchestrated by a B cell–lymphotoxin-dependent pathway. Blood, 115(23):4725–4733, June 2010. ISSN 0006-4971. doi: 10.1182/blood-2009-10-250118.

45. Inken D. Kelch, Gib Bogle, Gregory B. Sands, Anthony R. J. Phillips, Ian J. LeGrice, and P. Rod Dunbar. High-resolution 3D imaging and topological mapping of the lymph node conduit system. PLoS Biology, 17(12):e3000486, December 2019. ISSN 1544-9173. doi: 10.1371/journal.pbio.3000486.

46. Madeline A. Kibler, Margaux D. Miller, Michael J. Donzanti, and Jason P. Gleghorn. Confocal-Compatible Workflow for Sectioning, Staining, and Imaging Serial Vibratome Sections for 3D Anatomical Reconstruction of the Lymph Node, November 2025. ISSN: 2692-8205 Pages: 2025.10.29.685350 Section: New Results.

47. Luke Xie (2024). Hysteresis thresholding for 3D images (or 2D) (https://www.mathworks.com/matlabcentral/fileexchange/44648-hysteresis-thresholding-for-3d-images-or-2d), MATLAB Central File Exchange. Retrieved June 9, 2024.

48. Aaron Carass, Snehashis Roy, Adrian Gherman, Jacob C Reinhold, Andrew Jesson, Tal Arbel, Oskar Maier, Heinz Handels, Mohsen Ghafoorian, Bram Platel, and others. Evaluating white matter lesion segmentations with refined Sørensen-Dice analysis. Scientific reports, 10(1):8242, 2020. Publisher: Nature Publishing Group UK London.

49. Raneem Ismail and Szilvia Nagy. Ways of improving of active contour methods in colonoscopy image segmentation. Image Analysis and Stereology, 41(1), 2022.

50. Bowen Cheng, Ross Girshick, Piotr Dollár, Alexander C. Berg, and Alexander Kirillov. Boundary IoU: Improving Object-Centric Image Segmentation Evaluation, March 2021. arXiv:2103.16562 [cs].

51. Rachel M. Gilbert, Laurel E. Schappell, and Jason P. Gleghorn. Defective mesothelium and limited physical space are drivers of dysregulated lung development in a genetic model of congenital diaphragmatic hernia. 148(10): dev199460. ISSN 0950-1991. doi: 10.1242/dev.199460.

52. Jeffery J. Ballyns, Jason P. Gleghorn, Vicki Niebrzydowski, Jeremy J. Rawlinson, Hollis G. Potter, Suzanne A. Maher, Timothy M. Wright, and Lawrence J. Bonassar. Image-guided tissue engineering of anatomically shaped implants via MRI and micro-CT using injection molding. 14(7):1195–1202. ISSN 1937-3341. doi: 10.1089/ten.tea.2007.0186.

53. Brea Chernokal, Cailin R. Gonyea, and Jason P. Gleghorn. Lung development in a dish: Models to interrogate the cellular niche and the role of mechanical forces in development. In Chelsea M. Magin, editor, Engineering Translational Models of Lung Homeostasis and Disease, Advances in Experimental Medicine and Biology, pages 29–48. Springer International Publishing,. ISBN 978-3-031-26625-6. doi: 10.1007/978-3-031-26625-6_3.

54. Michael J. Donzanti, Bryan J. Ferrick, Omkar Mhatre, Brea Chernokal, Diana C. Renteria, and Jason P. Gleghorn. Stochastic to deterministic: A straightforward approach to create serially perfusable multiscale capillary beds. 11(1):239–248. doi: 10.1021/acsbiomaterials.4c01247. Publisher: American Chemical Society.

55. Joshua T. Morgan, Jasmine Shirazi, Erica M. Comber, Christian Eschenburg, and Jason P. Gleghorn. Fabrication of centimeter-scale and geometrically arbitrary vascular networks using in vitro self-assembly. 189:37–47,. ISSN 01429612. doi: 10.1016/j.biomaterials.2018.10.021.

56. Jason P. Gleghorn, Jiyong Kwak, Amira L. Pavlovich, and Celeste M. Nelson. Inhibitory morphogens and monopodial branching of the embryonic chicken lung - gleghorn - 2012 - developmental dynamics - wiley online library. 241(5): 852–862,.

57. Jason P. Gleghorn, Sriram Manivannan, and Celeste M. Nelson. Quantitative approaches to uncover physical mechanisms of tissue morphogenesis. 24(5): 954–961,. ISSN 1879-0429. doi: 10.1016/j.copbio.2013.04.006.

58. Kara E. Garcia, Wade G. Stewart, M. Gabriela Espinosa, Jason P. Gleghorn, and Larry A. Taber. Molecular and mechanical signals determine morphogenesis of the cerebral hemispheres in the chicken embryo. 146(20):dev174318. ISSN 1477-9129, 0950-1991. doi: 10.1242/dev.174318.

59. Isabelle Mondor, Audrey Jorquera, Cynthia Sene, Sahil Adriouch, Ralf Heinrich Adams, Bin Zhou, Stephan Wienert, Frederick Klauschen, and Marc Bajénoff. Clonal Proliferation and Stochastic Pruning Orchestrate Lymph Node Vasculature Remodeling. Immunity, 45(4):877–888, October 2016. ISSN 1097-4180. doi: 10.1016/j.immuni.2016.09.017.

60. A Jagarapu, MJ Piovoso, and R Zurakowski. An Integrated Spatial Dynamics—Pharmacokinetic Model Explaining Poor Penetration of Anti-retroviral Drugs in Lymph Nodes. Front. Bioeng. Biotechnol. 8: 667. doi: 10.3389/fbioe, 2020.

61. Cailin R. Gonyea, Yuanjun Shen, Katherine M. Nelson, Rylie N. Bird, Rachel M. Gilbert, Oluyinka O. Olutoye, Sundeep G. Keswani, and Jason P. Gleghorn. The nitrofen/bisdiamine murine model of congenital diaphragmatic hernia has a pulmonary hypertension vascular phenotype consistent with human CDH. ISSN 1522-1504. doi: 10.1152/ajplung.00233.2024.

62. Brea Chernokal, Bryan J. Ferrick, and Jason P. Gleghorn. Zonal patterning of extracellular matrix and stromal cell populations along a perfusable cellular microchannel. 24(23):5238–5250,. ISSN 1473-0189. doi: 10.1039/D4LC00579A. Publisher: The Royal Society of Chemistry.

63. Vani Narayanan, Laurel E. Schappell, Carl R. Mayer, Ashley A. Duke, Travis J. Armiger, Paul T. Arsenovic, Abhinav Mohan, Kris N. Dahl, Jason P. Gleghorn, and Daniel E. Conway. Osmotic gradients in epithelial acini increase mechanical tension across e-cadherin, drive morphogenesis, and maintain homeostasis. 30(4):624–633.e4. ISSN 1879-0445. doi: 10.1016/j.cub.2019.12.025.

64. Vonetta L Edwards, Elias McComb, Jason P Gleghorn, Larry Forney, Patrik M Bavoil, and Jacques Ravel. Three-dimensional models of the cervicovaginal epithelia to study host–microbiome interactions and sexually transmitted infections. 80(1):ftac026. ISSN 2049-632X. doi: 10.1093/femspd/ftac026.

